# Enhanced Processivity and Collective Force Production of Kinesin-1 at Low Radial Forces

**DOI:** 10.1101/2025.08.27.672644

**Authors:** Andrew M. Hensley, Ahmet Yildiz

**Affiliations:** Physics Department, University of California, Berkeley, CA, USA; Department of Molecular and Cell Biology, University of California, Berkeley, CA, USA

## Abstract

Kinesin-1 is a robust motor that carries intracellular cargos towards the plus ends of microtubules. However, optical trapping studies reported that kinesin-1 is a slippery motor that quickly detaches from the microtubule, and multiple kinesins are incapable of teaming up to generate large collective forces. This may be due to the vertical (z) forces that the motor experiences in a single bead trapping assay, accelerating the detachment of the motor from a microtubule. Here, we substantially lowered the z-force by using a long DNA handle between the motor and the trapped bead and characterized the motility and force generation of single and multiple kinesin-1s. Contrary to previous views, we show that kinesin-1 is a robust motor that resists microtubule detachment before it reaches high hindering forces, but it quickly detaches under assisting forces even at low z-forces. We also demonstrate highly efficient collective force generation by multiple kinesin-1 motors. These results provide an explanation for how multiple kinesins team up to perform cellular functions that require higher forces than a single motor can bear.

## Introduction

Cytoskeletal motors generate forces to perform a variety of functions in cells, such as anchoring organelles to the cytoskeleton, maintaining tension on the mitotic spindle, and transporting cargoes over long distances. Kinesin-1 (hereafter kinesin) is the primary motor involved in cargo transport towards the plus-ends of microtubules in all eukaryotes (Yildiz, 2024). The force-generating properties of kinesin have been studied extensively by optical trapping (Andreasson et al., 2015; Carter & Cross, 2005; Milic et al., 2014; Nishiyama et al., 2002; Svoboda & Block, 1994; Yildiz et al., 2008). In these assays, a motor carries an optically trapped bead on a surface-immobilized microtubule, and a stationary trap exerts a springlike force on the bead proportional to its displacement from the trap center. Eventually, the motor experiences a hindering force that renders it unable to move forward, referred to as the stall force. Optical trapping can also be used for force-clamping to monitor motor movement under constant forces (Visscher et al., 1999). Previous studies reported that kinesin stalls under 5-7 pN hindering forces and exhibits an asymmetric response to hindering and assisting forces (Andreasson et al., 2015; Carter & Cross, 2005; Dogan et al., 2015; Nishiyama et al., 2002; Svoboda & Block, 1994). Under increasing hindering forces, velocity and run length gradually approach zero at the stall force, while the detachment rate continues to increase beyond the stall force (Andreasson et al., 2015). In comparison, kinesin is far more sensitive to assisting forces, quickly detaching from the microtubule at lower forces (Andreasson et al., 2015). The velocity of the motor does not significantly increase under assisting forces, consistent with the view that kinesin stepping is not gated by the detachment of the trailing motor domain from the microtubule (Hancock, 2016).

Optical trapping studies showed that most processive runs of beads driven by single kinesin motors end abruptly and prematurely before the motor reaches its stall force (Pyrpassopoulos et al., 2020; Svoboda & Block, 1994; Uemura & Ishiwata, 2003). Additionally, collective force generation by multiple kinesins is not proportional to the kinesin copy number, with teams of kinesins appearing to generate only slightly more force than one motor (Elshenawy et al., 2019; Furuta et al., 2013; Jamison et al., 2012). This has been attributed to the slipping of kinesin motors from the microtubule under force (Sudhakar et al., 2021; Toleikis et al., 2020). According to these measurements, a cargo containing teams of competing kinesins and dyneins would be dominated by dynein, as the greater collective force generated by dynein would induce kinesin detachment (Elshenawy et al., 2019). However, in vitro assays that monitored tug-of-war between kinesin and dynein on artificial scaffolds have shown kinesin to be a strong competitor to active mammalian dynein (Belyy et al., 2016; Derr et al., 2012; Gicking et al., 2022). It remains unclear how kinesin can compete against dynein while quickly falling off the microtubule.

Recent experimental and theoretical studies showed that in vitro optical trapping measurements of kinesin are sensitive to the experimental geometry and the size of the trapped bead (500 to 1,000 nm in diameter), typically an order of magnitude larger than the motor (Khataee & Howard, 2019). Because the bead stays on top of a surface-immobilized microtubule, the tether that a motor forms between the bead and the microtubule is oriented at ∼70° angle relative to the microtubule (Fig. 1A). When the bead is being moved horizontally by the trap, the tethered motor is being pulled towards the center of the bead. The magnitude of the z-force is estimated to be several-fold larger than the horizontal force, and the direction of this force is always pointed away from the microtubule when the trap pulls the bead in assisting or hindering directions (Khataee & Howard, 2019; Pyrpassopoulos et al., 2020). In cells, motors are expected to experience different z-forces depending on the size and flexibility of intracellular cargos. For example, a motor carrying a small (∼50 nm diameter) vesicle experiences little to no z-force, while it may be subjected to large z forces when carrying larger cargos, such as mitochondria. In addition, phospholipid membranes are deformable under force (D’Souza et al., 2023), and kinesin is capable of pulling narrow membrane tethers (Leduc et al., 2004), which may reduce the magnitude of z-forces kinesin bears during intracellular transport.

**Figure 1.**
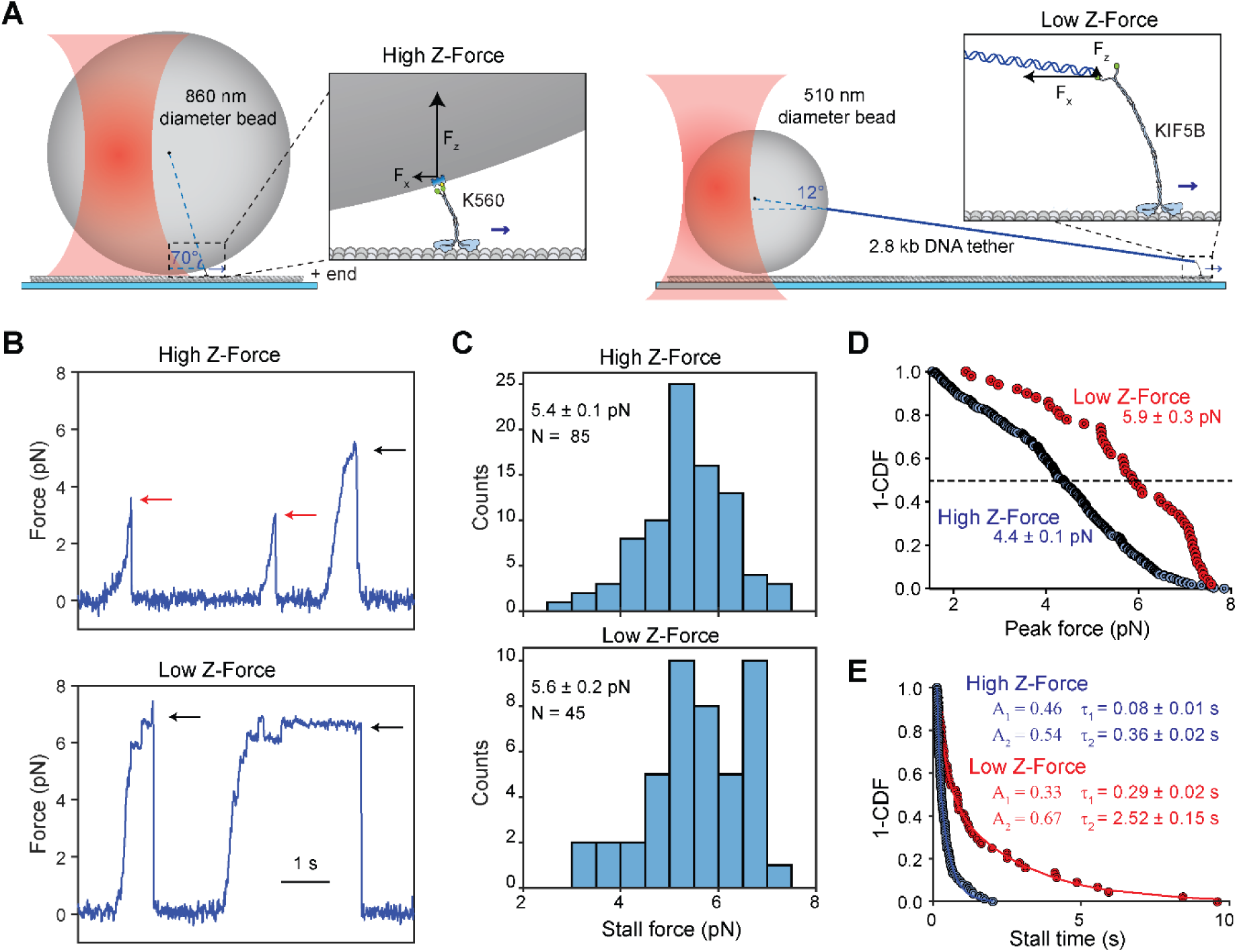
The DNA tethered bead assay to reduce the z-force in the optical trap. **A)** (Left) Conventional single-bead trapping assay where a truncated, biotinylated kinesin is attached to an 860-nm-diameter streptavidin-coated bead. (Right) The DNA-tethered bead assay, where full-length KIF5B is attached to a 510-nm-diameter bead via a 2800 bp DNA handle. **B)** Characteristic force-generating behavior of kinesin against a stationary trap in both a high and low z-force regime. Stalls are marked with black arrows, while premature detachments are marked with red arrows. **C)** Stall force of kinesin (mean ± s.e.m.) in the high and low z-force regimes. **D)** The inverse cumulative distribution function (1-CDF) of the peak forces (median ± s.e.m.) of every processive run that exceeded 1.5 pN hindering force in the high (N = 264) and low (N = 51) z-force regimes. **E)** 1-CDF of motor stall times. Solid curves represent fits to a double exponential decay to calculate the amplitude (A) and mean lifetime (τ; ± s.e.). The weighted averages of the two decays are 0.27 ± 0.02 s and 1.49 ± 0.08 s for high (N = 85) and low (N = 45) z-forces, respectively.

A recent study eliminated this z-force in a microtubule dumbbell assay where a microtubule is trapped with two beads attached on each end and lowered onto a third surface-immobilized bead sparsely decorated with kinesin (Pyrpassopoulos et al., 2020). In this assay, kinesin is subjected to a force that is nearly parallel to the microtubule, and unlike single-bead assays, most runs reach a full stall before the motor detaches from the microtubule. One limitation of this microtubule dumbbell assay is the potential for kinesin to be subjected to torsional constraints. This is because kinesin walks symmetric hand-over-hand (Yildiz et al., 2004), rotating its stalk counterclockwise 180° at every step (Ramaiya et al., 2017; Wolff et al., 2023). While a trapped bead is free to rotate around its center, a trapped microtubule is not, and this may result in warping of the coiled-coil stalk and hindering of kinesin motility (Ramaiya et al., 2017). Force exerted on the microtubule may also alter kinesin motility by altering the lattice stepping of microtubules (Memet et al., 2018; Shima et al., 2018). Another study connected kinesin to a microtubule via a long DNA tether and observed that a stalled motor detaches from the microtubule more slowly than a walking motor under unloaded conditions (Kuo et al., 2022). While this DNA tensiometer geometry fully eliminates the z-forces, unlike optical trapping, it is incapable of directly manipulating the motor by controlling the applied force.

In this study, we developed a single-bead trapping assay where kinesin is attached to a bead through a long double-stranded DNA tether (Urbanska et al., 2021). This assay geometry reduces the z-force on the motor by an order of magnitude and increases the degree of freedom of the motor attached to the bead, reducing the risk of torsional build-up during processive motility. In these low z-force assays, kinesin exhibited robust stalls and fewer premature detachments from the axoneme under hindering forces. We also observed higher asymmetry in the force-velocity relationship of kinesin under constant hindering forces, compared to conventional single-bead assays. Lowering of the z-force also enabled more efficient collective force generation by multiple kinesin motors carrying the bead. These results challenge the established views and show that kinesin is a robust motor that persists in microtubule detachment against large hindering forces and efficiently teams up for carrying force generation functions under high tension.

## Results

### Reducing the z-force increases the kinesin stall time

To investigate the role of z-forces in kinesin motility and force generation, we first performed fixed trapping assays using two different assay geometries. In a conventional single bead assay, we directly attached biotinylated kinesin motors to 860-nm-diameter streptavidin-coated polystyrene beads (Fig. 1A). We used a tail-truncated, constitutively active human kinesin-1 KIF5B (K560; amino acids 1-560; Figure 1_figure supplement 1A), as this is a commonly used construct in previous optical trapping studies of kinesin (Andreasson et al., 2015; Carter & Cross, 2005; Milic et al., 2014; Nishiyama et al., 2002; Svoboda & Block, 1994; Yildiz et al., 2008). In our revised DNA-tethered-bead assay, we used an oligo-labeled full-length kinesin (KIF5B, Figure 1_figure supplement 1A) attached to a smaller diameter (510 nm) bead through a long (2.8 kb) DNA handle (Fig. 1A and Figure 1_figure supplement 1A-B). Full-length kinesin was chosen for the reduced z-force assay because it has a longer and more flexible stalk than K560 and better represents the force behavior of kinesin under physiological conditions. Although full-length kinesin is mostly autoinhibited (Yildiz, 2024), the attachment of kinesin to the bead from its tail and the force exerted by the trap activate the motor for processive motility (Belyy et al., 2016; Coy et al., 1999; Svoboda & Block, 1994). The DNA handle was functionalized with biotin at one end and a single-stranded overhang on the other end. Kinesin was labeled with a complementary oligo at its C-terminus and attached to the handle through DNA hybridization (Figure 1_figure supplement 1C-E). This revised geometry results in an approximately 10-fold reduction in the z-force relative to the force parallel to the microtubule surface (from ∼2.5 Fx to ∼0.25 Fx, Fig. 1A).

Conventional trapping assays (high z-force) yielded results comparable to previous reports (Carter & Cross, 2005; Pyrpassopoulos et al., 2020; Svoboda & Block, 1994), where kinesin exhibited brief stalls at 5.4 ± 0.1 pN (mean ± s.e.m.) and most runs ended prematurely with rapid detachment from the microtubule (Fig. 1B-C). The stall forces collected under low z-force conditions were not significantly different (5.6 ± 0.2 pN; Student’s t-test, p = 0.16; Fig. 1B-C). To quantify the rate of premature detachment, we measured the peak force of every event exceeding 1.5 pN (Fig. 1D, see Methods). The median peak force in the low z-force regime was similar to the stall force (5.9 ± 0.3 vs 5.6 ± 0.2 pN), but the peak force in the high z-force regime was reduced compared to the corresponding stall force (4.4 ± 0.1 vs. 5.4 ± 0.1 pN). When peaks not corresponding to stalls were counted as premature detachments, only one third of the runs reached a stall in the high z-force regime, whereas almost 90% of the runs ended with a stalling event in the low z-force regime. Collectively, these results indicate that the z-forces lead to reduced stall times and increased frequency of premature detachment.

In addition to the higher probability of reaching a stall, durations at stall were longer when the z-force was minimized. The low z-force assay produced stalls with visibly longer durations (Fig. 1B, p < 0.0001). 40% (18 out of 45) of stalls lasted more than 1 s in the low z-force regime compared to less than 10% (7 out of 85) in the high z-force regime (Fig. 1E). Stall times were fit to a double exponential decay (Fig. 1E), producing time constants that were 3- to 7-fold higher in the low z-force regime compared to the high z-force regime. These time constants are comparable to those reported for minus-end-directed dynein under high z-forces (Elshenawy et al., 2020). We also noticed that the average stall duration of kinesin under low z force (1.49 s) was slightly shorter than the run time of kinesin in unloaded conditions (6.17 s, Figure 1_figure supplement 1F-G), suggesting that hindering forces do not increase the time kinesin remains bound to the microtubule. Our average stall duration is comparable to those recently reported by the three bead trapping assays (1.3 s) (Pyrpassopoulos et al., 2020), but shorter than the DNA tensiometer assays (3 s) (Noell et al., 2024). This discrepancy may be attributed to the presence of low z forces in our trapping assays that increase the rate of detachment, compared to the nearly parallel force motor experiences in DNA tensiometer assays. In addition, kinesin may be more likely to fully disengage from the microtubule and return to the trap center in optical trapping assays, whereas it can more frequently exhibit short backward slips and keep walking on the microtubule in the DNA tensiometer assay (Noell et al., 2024), resulting in longer apparent stall durations.

### Kinesin detachment rate is more asymmetric in the low z-force

Optical trapping studies revealed that kinesin rapidly releases from microtubules when pulled forward, while resisting backward movement when pulled towards the minus-end (Andreasson et al., 2015). A recent theoretical study predicted that the detachment rate of kinesin increases with assisting force, whereas it decreases with hindering force due to catch bond behavior (Khataee & Howard, 2019). The z-force is proportional to the parallel force exerted on the motor, and it results in a similar increase in the detachment rate under the same magnitude of hindering and assisting forces, reducing the overall asymmetry in force-detachment of kinesin (Khataee & Howard, 2019). Therefore, kinesin is expected to exhibit even greater asymmetry in its force-detachment kinetics in the absence of z-forces.

To test this model, we performed force feedback measurements on single kinesin motors in both the high and low z-force regimes at a variety of hindering and assisting forces (Fig. 2A, Figure 2_figure supplement 1). Under hindering forces, kinesin walked 2- to 4-fold farther at low z-forces (Fig. 2B, Figure 2_figure supplement 2A-B). Processive motility near the stall force (6 pN) was negligible, and we did not observe processive backward runs under superstall forces (10 pN), as previously reported (Figure 2_figure supplement 2C-D) (Andreasson et al., 2015). Detachment rates under hindering forces were uniformly lower in low z-force compared to high z-force, but the rate increased with the magnitude of force under both conditions. Under assisting forces, run lengths were shorter and detachment rates were higher than those measured under hindering forces (Andreasson et al., 2015), and we did not observe major differences between high and low z-force conditions (Fig. 2B, Figure 2_figure supplement 2A-B). We concluded that kinesin forms an asymmetric slip bond under horizontal forces, and microtubule detachment is more sensitive to assisting forces than hindering forces. This is consistent with the model that assisting forces result in a conformation that facilitates the release of Pi, resulting in the rapid detachment of the tethered head in a weakly bound ADP state (Hancock, 2016; Kawaguchi & Ishiwata, 2001; Milic et al., 2014), on a timescale faster than z-force-induced detachment. Therefore, reduction of the z-force in a DNA tethered trapping assay does not result in a substantial increase in motor run length under assisting forces.

**Figure 2.**
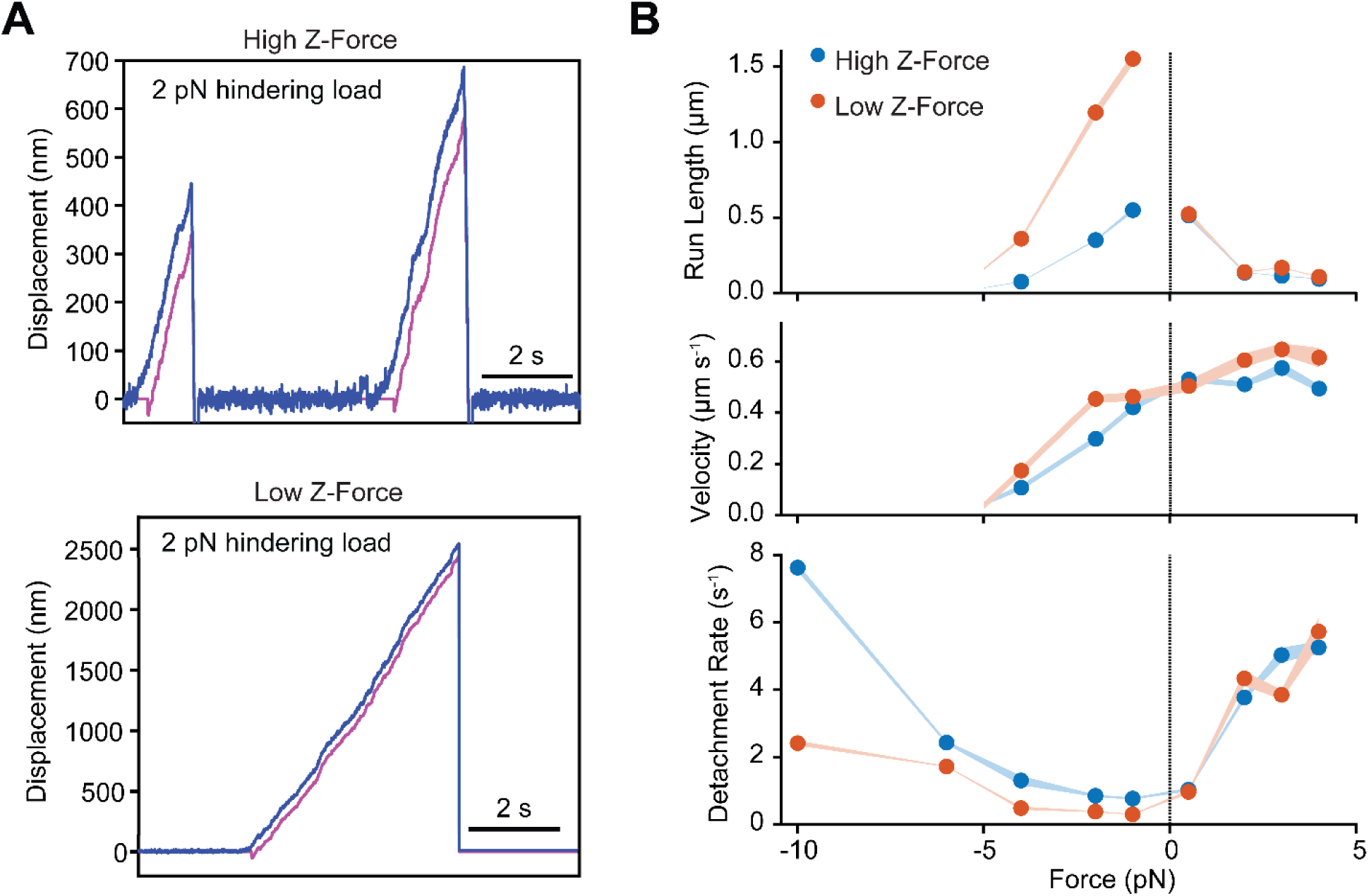
Kinesin exhibits asymmetric slip bond behavior under force. **A)** Example traces of kinesin motility under a 2 pN hindering force in high and low z-forces. The trap position (magenta) is updated to remain 100 nm behind the bead position along the microtubule long axis (blue). The trap stiffness is fixed at 0.02 pN nm^-1^ so that this separation corresponds to 2 pN. **B)** Run lengths (fit parameter from single-exponential CDFs, ± s.e.), velocities (mean ± s.e.), and detachment rates (defined as the ratio of run length to velocity of the motor) for forces ranging from –4 pN to +4 pN under high (from left to right, N = 60, 69, 65, 50, 167, 100, 115 runs) and low (from left to right, N = 89, 76, 44, 42, 65, 32, 39 runs) z-force conditions. For –10 pN and –6 pN hindering forces, detachment rates were calculated from the run time distributions under high (N = 133, 167) and low (N = 43, 73) z-force regimes (Figure 2_figure supplement 2), as these conditions yielded minimal forward displacement. Negative and positive forces correspond to hindering (minus-ended) and assisting (plus-ended) directions, respectively. Under hindering forces, kinesin runs several-fold longer distances and has a lower detachment rate in the low z-force regime compared to the high z-force regime.

We also noticed that kinesin walks slightly faster in DNA-tethered bead assays (Fig. 2B). This could be due to partial hindrance of kinesin motility when it is directly linked to a large bead. Although a spherical bead is expected to be free to rotate as kinesin walks, its large torsional stiffness may result in supertwisting of the kinesin motor (Ramaiya et al., 2017), resulting in slower motility. In the DNA-tethered bead assay, rotational motion of kinesin may be distributed along the length of the DNA handle and the longer and more flexible stalk of full-length kinesin, which may mitigate this effect.

### Teams of kinesin generate more force under reduced radial force conditions

Previous optical trapping studies have attempted to characterize the force generated by teams of kinesin under single-bead optical trapping (Bormuth et al., 2008; Elshenawy et al., 2019; Furuta et al., 2013; Jamison et al., 2012). Unlike dyneins (Elshenawy et al., 2019), kinesins are not able to generate substantially higher forces than a single motor when a known number of motors are assembled on DNA origami and pulled by an optical trap (Elshenawy et al., 2019; Furuta et al., 2013; Jamison et al., 2012). This is in contrast to the behavior of kinesin in vivo, where two kinesins can generate more force than a single motor (Shubeita et al., 2008). In addition, multiple surface-immobilized kinesin motors were shown to glide microtubules against much larger collective forces than a single motor can produce (Bormuth et al., 2008; Kuo et al., 2022). One potential explanation for this discrepancy is that the premature detachment of kinesin from microtubules under high z-forces limits the number of kinesins that can be engaged simultaneously with the microtubule in these assays.

We tested whether teams of kinesins could generate more collective force under low z-force conditions than under the high z-force conditions. For the high z-force conditions, we created a short DNA chassis that recruits either two or three K560 motors directly to a bead (Fig. 3A, Figure 3_figure supplement 1A). For the low z-force condition, we used a modified chassis that binds to a long DNA handle on the bead via DNA hybridization (see Methods, Fig. 3A-B, Figure 3_figure supplement 1B). In multiple motor assays, we used constitutively active K560 as a motor in both conditions to eliminate the possibility of autoinhibition of kinesin bound to the DNA chassis (Coy et al., 1999; Friedman & Vale, 1999). To estimate the percentage of chassis with two and three motors bound, we performed mass photometry measurements at a 3-fold molar excess of kinesin to the chassis, as higher ratios would obscure the distinction of complexes from the kinesin-only population. Assuming there is no cooperativity among the binding sites, we modeled motor occupancy using a Binomial distribution (Figure 3_figure supplement 2). We observed 17-29% of particles corresponded to the two-motor species on the 2-motor chassis in mass photometry, indicating that 45-78% of the 2-motor chassis was bound to two kinesins. Similarly, 15% and 40% of the 3-motor chassis were bound to two and three kinesins, respectively.

**Figure 3.**
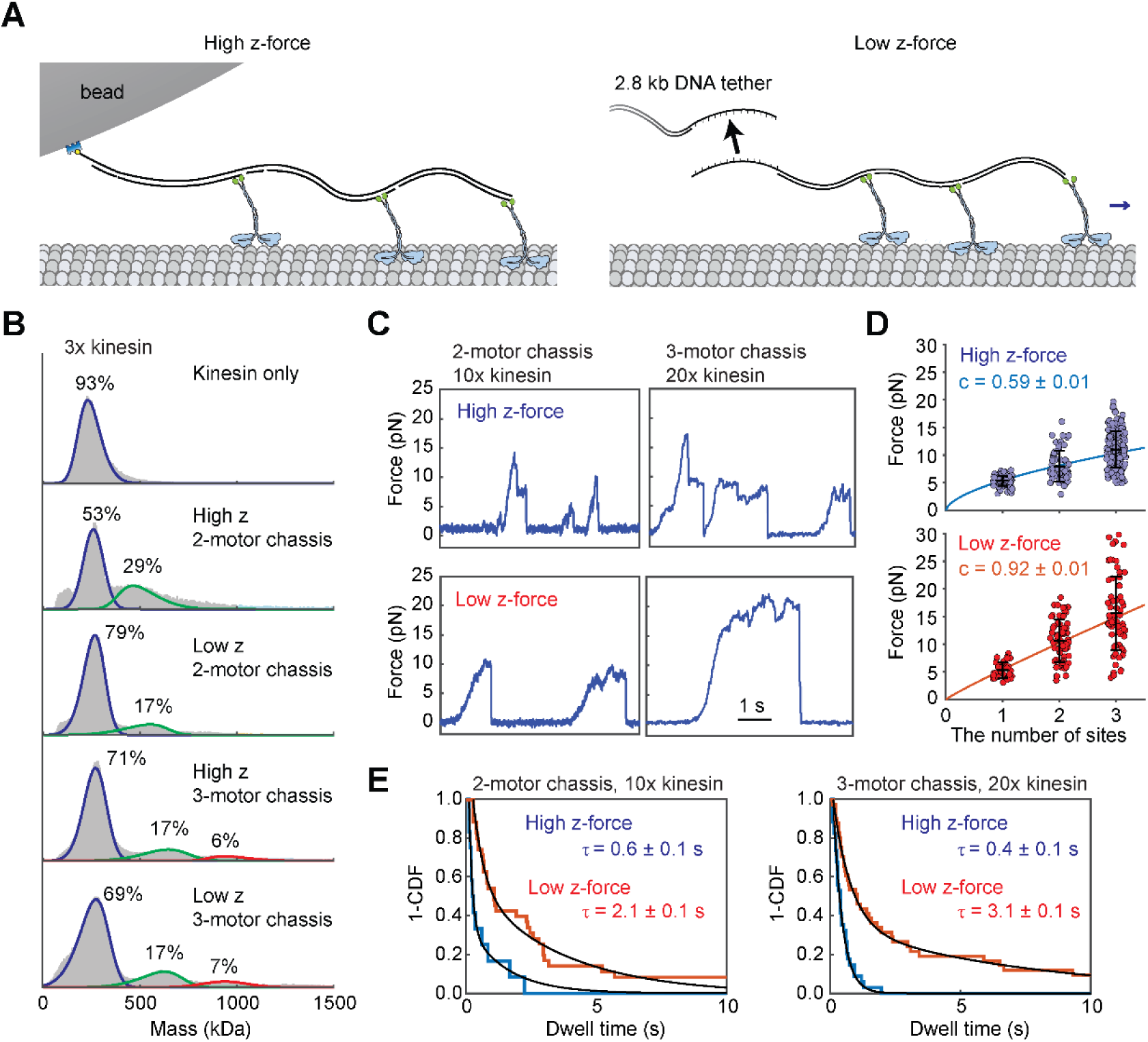
Kinesin motors team up efficiently under low z-force conditions. **A)** Schematic of the high and low z-force assays with three K560 motors bound to a DNA chassis. Left: The 3-motor DNA chassis binds directly to a streptavidin-coated bead. Right: The DNA chassis with a single-stranded DNA overhang hybridizes to the overhang on the long DNA handle attached to the bead. **B)** Mass photometry measurements with a DNA chassis are performed with a 3-fold excess of kinesin. The Gaussian mixture model identifies the mass and percentage of unbound kinesin (blue curves), chassis with one (blue curves), two (green curves), and three (red curves) motors. Mass populations of unbound motor and chassis with one motor bound cannot be distinguished from each other. **C)** Example trajectories of beads driven by two and three K560 motors in the high and low z-force regimes. These assays were performed with a 10 or 20-fold excess of kinesin for 2- or 3-motor DNA chassis, respectively. **D)** Forces (mean ± s.d.) produced by single kinesins or kinesins bound to 2- or 3-motor chassis under high (from left to right, N = 85, 66, 158) and low z-force conditions (from left to right, N = 37, 65, 66). The solid line represents a fit to a power function to calculate the scaling factor (c, ± s.e.). **E)** Dwell times before slips are plotted as 1-CDF. Weighted average time constants (τ, ± s.e.) are extracted from fitting to a double exponential decay.

In optical trapping assays, we used 10-fold and 20-fold molar excess of kinesin for 2-motor and 3-motor chassis, respectively, to substantially increase the percentage of the chassis carried by multiple kinesins. Under these conditions, we estimate 76-93% of the 2-motor chassis were bound to two kinesins, and 30% and 70% of 3-motor chassis were bound to two and three kinesins, respectively. Chassis with a single kinesin was predicted to be negligible under these conditions. In the high z-force condition, 2- and 3-motor chassis were able to generate 8.0 ± 0.3 pN and 11.0 ± 0.3 pN, respectively. By comparison, in the low z-force condition, 2- and 3-motor chassis were able to generate 10.6 ± 0.5 pN and 15.6 ± 0.8 pN (Fig. 3C-D, Figure 3_figure supplement 3), respectively. We computed a scaling factor by fitting the average measured forces to a power equation *F = F_s_ · x^c^* where *F̄* is the average measured force, 𝐹_𝑆_ is the stall force of a single kinesin, 𝑥 is the number of motor binding sites on the chassis, and 𝑐 is the scaling factor. Consistent with previous reports (Elshenawy et al., 2019; Furuta et al., 2013; Jamison et al., 2012), we observed a subproportional (𝑐 = 0.59 ± 0.01) increase in stall force by the motor copy number in the high z-force condition. In the low z-force condition, 𝑐 was 0.92 ± 0.01, indicating nearly proportional force generation (Fig. 3D).

We also tested how z-forces affect the backward slipping of kinesin-driven beads without fully disengaging from the microtubule (Elshenawy et al., 2019; Furuta et al., 2013; Jamison et al., 2012). Here, we define a slip as an abrupt backward movement that decreases the force on the bead by more than 2 pN, followed by switching back to a motility-competent state without returning to the trap center. Remarkably, we observed beads driven by teams of kinesins slipping from the microtubule less frequently in the low z-force condition (Fig. 3C,E). In comparison, slipping events were rarely observed in beads driven by a single motor, suggesting that kinesin typically detaches rather than slipping back on the microtubule under hindering loads.

Slippage of beads driven by multiple motors may correspond to the microtubule detachment of the leading motor while the trailing motors remain attached, which would result in backward movement of the bead without disengaging it from the microtubule (Jamison et al., 2012). Beads driven by single kinesins were also reported to slip backward under hindering forces, without disengagement of the motor from the microtubule(Sudhakar et al., 2021; Toleikis et al., 2020). Our results indicate that teams of kinesins are capable of reaching and sustaining motility under high hindering forces, whereas their collective force generation is limited by premature detachment or slippage of individual motors under high z-forces exerted by an optical trap. In comparison, kinesins persistently bind to the microtubule and resist slipping under low z-forces, which increases their efficiency to perform work against superstall forces.

## Discussion

In this study, we used a long DNA handle to substantially reduce the z-force experienced by kinesin in a single-bead optical trapping assay (Urbanska et al., 2021). Our work shows that kinesin is sensitive to z-forces, which causes the motor to reach a stall less frequently and stall for shorter durations. Reducing the z-force enhances the processivity and slows detachment under constant hindering forces (Fig. 4A) (Urbanska et al., 2021), but not under assisting forces. Our results agree with the theoretical prediction that kinesin exhibits higher asymmetry in force-detachment kinetics without z-forces (Khataee & Howard, 2019), and are consistent with optical trapping and DNA tensiometer assays that reported more persistent stalling of kinesin in the absence of z-forces (Kuo et al., 2022; Noell et al., 2024; Pyrpassopoulos et al., 2020).

**Figure 4.**
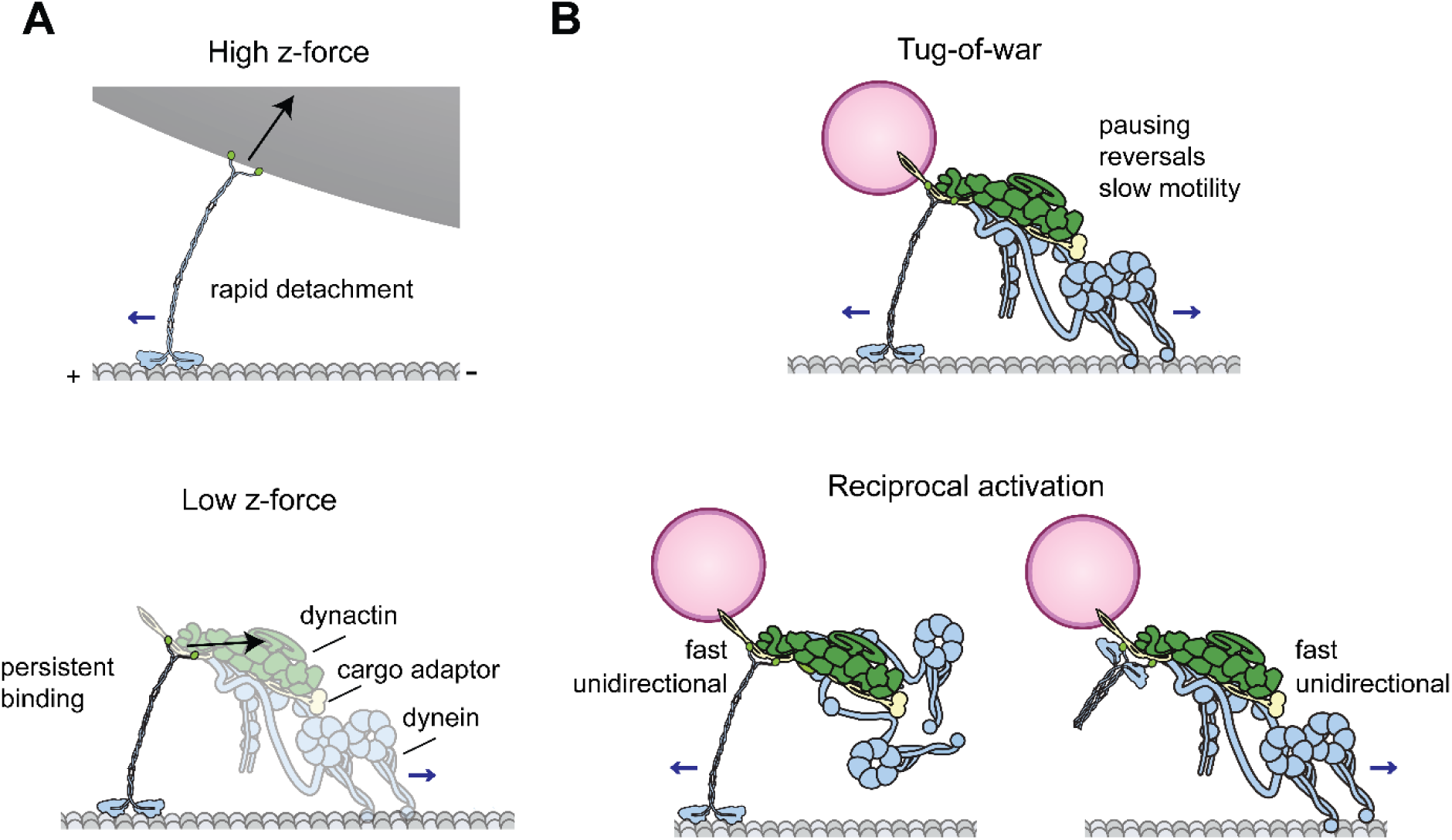
Kinesin resists microtubule detachment and backward processive movement under low z forces. **A)** High z-forces inherent to optical trapping assays result in rapid detachment of kinesin from microtubules, whereas the motor experiences lower z-forces when carrying intracellular cargos and more persistently binds to microtubules under those conditions. **B)** (Top) The tug-of-war between kinesin and dynein results in pausing or stalling the cargo movement, as well as slow motility interspersed with frequent reversals, because either motor resists detachment or backward stepping under forces generated by its opponent. (Bottom) Coordination between the opposing motors, such as activation of one motor at a time (reciprocal activation) results in fast unidirectional transport in the direction of an active motor while the inactive motor is carried as a passive cargo.

Force-detachment kinetics of protein-protein interactions have been modeled as either a slip, ideal, or catch bond, which exhibit an increase, no change, or a decrease in detachment rate, respectively, under increasing force (Thomas et al., 2008). Slip bonds are most commonly observed in biomolecules, but studies on cell adhesion proteins reported a catch bond behavior (Marshall et al., 2003). Although previous trapping studies of kinesin reported a slip bond behavior (Andreasson et al., 2015; Carter & Cross, 2005), recent DNA tensiometer studies that eliminated the z-force showed that the detachment rate of the motor under hindering forces is lower than that of an unloaded motor walking on the microtubule (Kuo et al., 2022; Noell et al., 2024), consistent with the catch bond behavior. Unlike these reports, we observed that the stall duration of kinesin is shorter than the motor run time under unloaded conditions, and the detachment rate of kinesin increases with the magnitude of the hindering force. Therefore, our results are more consistent with the asymmetric slip bond behavior. The difference between our results and the DNA tensiometer assays (Kuo et al., 2022; Noell et al., 2024) can be attributed to the presence of low z-forces in our DNA-tethered optical trapping assays, which may increase the detachment rate under high hindering forces. Future studies that could directly control hindering forces and measure the motor detachment rate in the absence of z-forces would be required to conclusively reveal the bond characteristics of kinesin under hindering loads.

It is also possible that the hindering force slows microtubule detachment of kinesin by trapping the motor in a high microtubule affinity state during stepping (Noell et al., 2024), rather than increasing the bond strength between kinesin and the microtubule. Consistent with this view, hindering force was shown to stabilize dynein’s coiled-coil stalk in a registry that favors high affinity for the microtubule (Rao et al., 2019), and to trap myosin I in an ADP-bound state that strongly binds to actin (Guo & Guilford, 2006; Laakso et al., 2008). A similar mechanism may favor more persistent microtubule binding of kinesin under hindering forces, even though the force induces faster detachment from the microtubule (Noell et al., 2024).

Rapid detachment induced by z-forces has implications for the way teams of kinesin cooperate to generate forces collectively. Premature detachment of kinesins under low hindering forces hampers collective motility against larger hindering forces by limiting the number of kinesins simultaneously engaging with the microtubule (Ucar & Lipowsky, 2020). We observed that a team of kinesins frequently undergoes sequential slips when it reaches large hindering forces under high z-force conditions (Fig. 3C, E). However, under low z-force conditions, collective force generation was nearly proportional to the motor copy number. These results explain why previous single-bead trapping measurements under high z-forces failed to observe the collective force generation of multiple kinesins (Elshenawy et al., 2019; Furuta et al., 2013; Jamison et al., 2012) and reconcile the results of these measurements with the ability of multiple kinesins to work against high hindering forces under different experimental geometries (Bormuth et al., 2008; Kuo et al., 2022).

Our results also have implications for how kinesin can function together with dynein to drive bidirectional cargo transport. The tug-of-war model suggests opposing teams of dyneins and kinesins undergo mechanical competition to determine which direction their cargo moves (Fig. 4B) (Hancock, 2014). Attempts to study the tug-of-war model in vitro have yielded mixed results. Kinesin was previously shown to be a weak competitor to yeast dynein (Derr et al., 2012), while other studies have shown kinesin to be competitive against mammalian dynein (Belyy et al., 2016; Gicking et al., 2022). Regardless of which motor wins the tug-of-war, these studies consistently reported slow unidirectional velocities well below those exhibited by single motors, and reversal of directionality was extremely rare (Belyy et al., 2016; Derr et al., 2012; Gicking et al., 2022). This contrasts with intracellular transport, where cargos typically move unidirectionally at nearly the full speed of the active motor and frequently stall and reverse their direction (Cason & Holzbaur, 2022). Computational studies predicted that the velocity and directionality of tug-of-war are highly sensitive to the detachment rate of the motors (Ma et al., 2023; Ohashi et al., 2019). These models predicted that motors that detach quickly from the microtubule are less likely to win the tug-of-war, even if they can walk against higher hindering forces. Unlike previous reports (Andreasson et al., 2015; Block et al., 2003), we showed that kinesin is not a slippery motor and exhibits a low detachment rate under hindering forces when the z-force inherent to optical trapping is minimized. These results are consistent with the observed slow motility when kinesin and dynein are pitted against each other in a tug-of-war (Derr et al., 2012; Elshenawy et al., 2019; Gicking et al., 2022). Moreover, we demonstrated that teams of kinesin can generate forces proportional to the motor copy number, similar to dynein (Elshenawy et al., 2019), indicating that multiple kinesins cannot be readily forced to move backwards under minus-end-directed forces generated by a team of dyneins on the same cargo.

Collectively, both kinesin and dynein exhibit highly asymmetric force detachment kinetics that favor microtubule detachment under assisting forces in their direction of motion, whereas they resist detachment under hindering forces (Andreasson et al., 2015; Dogan et al., 2015; Ezber et al., 2020; Pyrpassopoulos et al., 2020; Rao et al., 2019). Therefore, the tug-of-war mechanism is inconsistent with the fast unidirectional movement of intracellular cargos. Instead, it may be more relevant to immobilization or stalling of a substantial fraction of organelles and vesicles in cells (Noell et al., 2024; van Spronsen et al., 2013; Wiemer et al., 1997) or transient pauses of the cargo before it continues to move in the same direction or reverses its direction.

Our results also indicate that unidirectional transport of the cargo at near the full speed of either anterograde or retrograde motor would require inactivation of the opposing motor on the cargo, but the underlying mechanism is still emerging. Recent studies have shown that kinesin and dynein can be simultaneously recruited to a cargo with the same cargo adaptor protein (Canty & Yildiz, 2020; Reck-Peterson et al., 2018; Splinter et al., 2012). In vitro reconstitution of bidirectional transport of these complexes hinted at various mechanisms that coordinate the activity of these opposing motors when they form a complex with their adaptor (Fig. 4B) (Abid Ali et al., 2025; Canty et al., 2023; Fenton et al., 2021; Fu & Holzbaur, 2013; Kendrick et al., 2019). Future studies will be required to dissect whether cargos avoid tug-of-war for fast and efficient transport while utilizing the tug-of-war mechanism for pausing and reversal of cargo movement.

## Materials and Methods

### Construction of a Long DNA Handle

A PCR reaction was performed with 2 μl of 500 μg/mL lambda phage DNA (NEB), 15 μl each of Bio Forward and Kinesin Reverse primers (Table 1), 2.5 mL of KOD Hot Start Master Mix (NEB), and nuclease-free water up to 5 mL. This volume was subdivided into 96 tubes, and the PCR reaction was performed with annealing the primers at 57 °C for 10 s and extension at 70 °C for 85 s for 40 cycles. The resulting product was cleaned up using a PCR purification kit (Qiagen) and subsequently digested with BstXI for 2 h at 37 °C followed by heat inactivation of the enzyme at 80 °C for 20 min. The DNA handle was run on a 0.8% TAE agarose gel and the digested band was recovered via gel extraction. Oligos Kinesin 24B Bottom and 45B Top (Table 1) were annealed at 95 °C for 2 min in annealing buffer (10 mM Tris, 50 mM NaCl, 5 mM EDTA, pH 8) and gradually cooled to room temperature over 1 h. The annealed oligos were then ligated onto the digested DNA handle by incubating with T4 polynucleotide kinase and T4 DNA ligase at ambient temperature overnight in the T4 DNA ligase reaction buffer. Excess oligo was removed with a PCR purification kit.

**Table 1.**
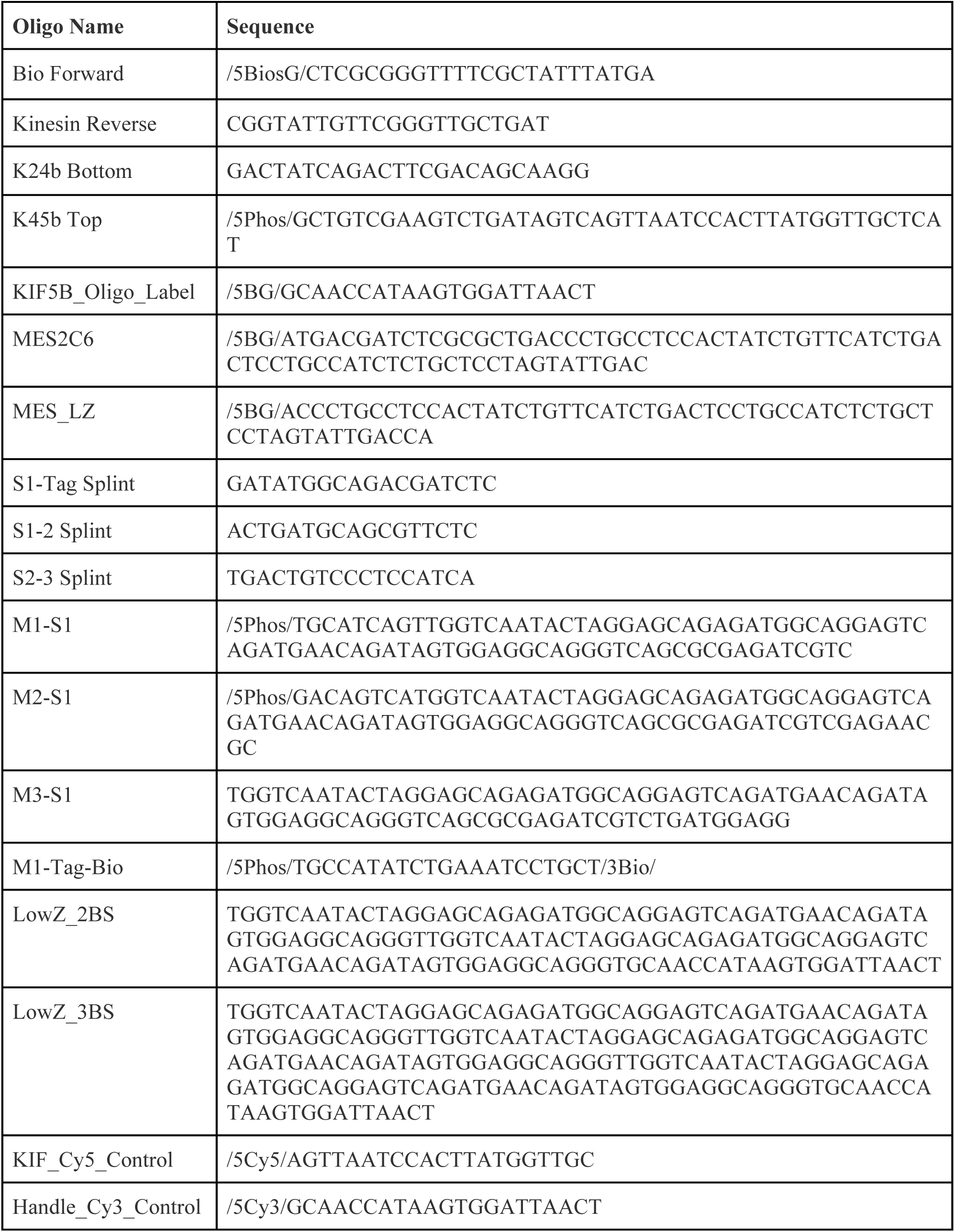
The list of DNA constructs used in this study.

The presence of the single-stranded overhang was confirmed by hybridizing the handle with excess Cy3-labeled complementary oligomers (Table 1) and running it on an agarose gel. The gel was subsequently imaged in the Cy3 channel of a Typhoon FLA 9500 (Cytiva) to verify colocalization of the Cy3 oligo with the handle (Figure 1_figure supplement 1B).

### Construction of the DNA Chassis

Four versions of the DNA chassis were used for multiple motor experiments. For the high z-force geometry, splint and backbone oligos were used as described previously (Elshenawy et al., 2019). The 2-motor chassis used the following oligos: M1-Tag-Bio, S1-Tag Splint, M1-S1, S1-2 Splint, M2-S1, whereas the 3-motor chassis used two additional oligos: S2-3 Splint and M3-S1 (Table 1). Briefly, 70 μl of biotinylated oligo was added to 50 μl backbone and 100 μl splint DNA oligos for the relevant constructs, annealed at 95 °C for 2 h, and gradually cooled to room temperature over 1 h. The resulting product was run on a 10% TBE gel (Cytiva). Bands containing the fully assembled chassis were extracted from the gel, flash frozen in liquid nitrogen, and crushed with a metal spatula. The crushed gel slices were then nutated in the crush and soak buffer (300 mM NaOAc, 1 mM EDTA, 0.1% SDS, pH 8) at room temperature for 48 h. After centrifugation at 5,000 g for 2 min, the supernatant was collected and passed through a DNA spin column (EconoSpin) to remove remaining polyacrylamide. The DNA was further purified using ethanol precipitation and resuspended in 10 mM Tris, pH 8. The concentration was determined from the absorbance at 260 nm using a Nanodrop 1000 (ThermoFisher). For reduced z-force experiments, long ultramer oligos were ordered from IDT (Table 1) and diluted in 10 mM Tris pH 8 for a final concentration of 100 μM.

### Purification and Labeling of Kinesin

Human full-length (ZZ-KIF5B-GFP-SNAP) and truncated (ZZ-KIF5B(1-560)-GFP-SNAP) kinesin-1 plasmids were encoded into a pOmnibac backbone and transformed into DH10Bac competent cells. The cells were plated onto bacmid plates at 37 °C for 48 h. A single white-colored colony was selected and grown overnight in 2X-YT media supplemented with 50 μg/mL kanamycin, 7 μg/mL gentamicin, and 10 μg/mL tetracycline. 2 mL of SF9 cells at 10^6^ cells/mL were adhered to one well in a 6-well TC plate. 192 μl of purified bacmid DNA at 1 μg/μL and 6 μl of FuGene HD transfection reagent were then mixed and added dropwise to the well. The plate was incubated at 27 °C for 72 h. 1 mL of the p1 virus was harvested from the well and used to infect 50 mL SF9 cells at 10^6^ cells/mL, which were then incubated at 27 °C for 72 h. Cells were centrifuged at 4,000 g for 10 min, and the supernatant collected as the p2 virus.

For protein expression, 10 mL of the p2 virus was added to 1 L of SF9 cells at 10^6^ cells/mL and incubated at 27 °C for 72 h. Cells were pelleted by centrifugation at 4000 *g* for 10 min. Pellets were washed with PBS buffer and either used immediately or flash frozen and stored at -80 °C. For protein purification, cells were first resuspended in lysis buffer (25 mM HEPES, 1 M KCl, 1 mM DTT, 1 mM PMSF, 0.1 mM ATP, one Roche protease inhibitor tablet per 100 mL, 10% glycerol, pH 7.4) and then lysed in a dounce homogenizer (Wheaton). The lysate was clarified by centrifuging at 65,000 rpm for 45 min in a Type 70 Ti rotor (Beckman Coulter). The lysate was then added to 1 mL IgG Sepharose 6 Fast Flow affinity resin (Cytiva) and rolled slowly for 2 h at 4 °C. The beads were then applied to a 20 mL gravity column (Bio-Rad) and subsequently washed with 4 column volumes lysis buffer and 1 column volume TEV buffer (25 mM HEPES, 300 mM KCl, 10 mM MgCl2, 1 mM EGTA, 1 mM DTT, 1 mM PMSF, 0.1 mM ATP, one Roche protease inhibitor tablet per 100 mL, 10% glycerol, pH 7.4).

Beads were then collected and labeled with 5 nmoles of custom benzylguanine (BG)-functionalized oligos (Biomer) or 2 nmoles of BG-Biotin (NEB) by rolling slowly overnight. Full-length KIF5B-GFP-SNAP was labeled with KIF5B_Oligo_Label (Table 1). For multi-motor assays, K560-GFP-SNAP was labeled with MES2C6 or MES_LZ for high or low z-force regimes, respectively (Table 1). Beads were applied to a 20 mL gravity column and washed with 5 column volumes of TEV buffer to remove excess oligo. Collected beads were incubated with 0.1 mg/mL TEV protease and nutated at room temperature for 1 hr. The beads were then applied to the gravity column, and the cleaved motor was collected from the flow through. This kinesin was then concentrated using a 100 kDa molecular weight cut-off (MWCO) centrifugal filter (Amicon) and its concentration determined by 280 nm absorbance using a Nanodrop 1000 (ThermoFisher). Aliquoted kinesin was flash frozen and stored at -80 °C.

All constructs were run on 4-12% Bis-Tris SDS-PAGE gels (Invitrogen) alongside a protein ladder (BIO-RAD Precision Plus). GFP- and SNAP-tagged full-length and truncated KIF5B heavy chains were clearly visible at their expected molecular weights (156 kDa and 110 kDa, respectively; Figure1_figure_supplement 1A). The DNA-labeled K560 was clearly separable from unlabeled K560 in an SDS-PAGE gel (Figure 3_figure supplement 1A), indicating a labeling efficiency of ∼50%. However, DNA-labeled KIF5B was not separable from unlabeled KIF5B in a gel, and its DNA labeling was confirmed by performing motility assays with a complementary Cy5 Oligo (Table 1, Figure 1_figure supplement 1C). For multi-motor experiments using a DNA chassis, 100 or 200 pmoles of oligo-labeled K560 was added to 10 pmoles of 2- or 3-motor chassis, respectively, and incubated overnight at 4 °C. The assembled complex was aliquoted, flash frozen, and stored at -80 °C the following day.

### Mass Photometry

Mass photometry measurements were acquired on a TwoMP mass photometer using six-well-cassette sample slides (Refeyn). 10 μl sample buffer (30 mM HEPES, 300 mM KOAc, 1 mM EGTA, 2 mM MgCl2, pH 7.4) was first added to each well for the droplet-dilution autofocus. Next, a mixture containing 100 nM K560 and 380 nM K560 was added to the well, and a 120-s movie was recorded with a large field of view. Mass measurements were calibrated using a standard mix composed of conalbumin, aldolase, and thyroglobulin. Mass photometry profiles were combined and fitted to multiple skewed Gaussian peaks, and their mean, standard deviation, and percentages were calculated using custom MATLAB software. Site occupancy of 2- and 3-motor chassis was estimated from the percentages of these populations using a Binomial model (Figure 3_figure supplement 2).

### Sample preparation for optical trapping assays

Flow chambers were built by sandwiching a glass slide and coverslip with permanent double-sided tape. When trapping single K560 motors, biotinylated K560 was added to 0.2% w/v 860-nm-diameter streptavidin-functionalized polystyrene beads (Spherotech) for 10 min. When trapping KIF5B, 2 μl of 100 nM DNA handle was incubated with 1 μl of 600 nM oligo-labeled KIF5B for 30 min on ice, before the mixture was diluted and added to 0.1% w/v 510-nm-diameter streptavidin-functionalized polystyrene beads (Bangs Labs) for 10 min. During this incubation time, axonemes purified from sea urchin sperm were diluted 1:10 in motor buffer (MB: 30 mM HEPES, 5 mM MgSO4, 1 mM EGTA, 10% glycerol, pH 7.4) for 3 min. The chamber was then washed with MBC (MB supplemented with 1.25 mg ml^−1^ casein and 1 mM DTT). Finally, the motor-bound beads were diluted 1:10 in stepping buffer (SB: MBC supplemented with 0.1 mg ml^−1^ glucose oxidase, 0.02 mg ml^−1^ catalase, 0.8% D-glucose, and 2 mM ATP).

Multi-motor trapping assays were performed similarly using 10x and 20x kinesin for 2- and 3-motor chassis, respectively. To estimate the percentage of chassis with multiple motors, we used the probability of kinesin binding to a site on a chassis from mass photometry in 3x excess condition to compute an effective dissociation constant 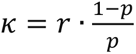, where *r* is the molar ratio of kinesin to chassis. Single-site occupancy at higher molar excesses of kinesin was calculated using this parameter.

In the high z-force regime, the biotinylated chassis was incubated with 0.2% w/v 860-nm-diameter polystyrene beads. In the low z-force regime, the chassis contains a single-stranded overhang complementary to the overhang of the DNA handle. 2 μl of 100 nM DNA handle was added to 1 μl of 600 nM K560-labeled chassis. Due to the ability of multiple kinesins to move against high hindering forces, 860 nm beads were used for the low z-force geometry to reduce the power of the trapping laser. For all multi-motor measurements, MBC and SB were additionally supplemented with 100 mM NaCl and 1 mg/mL BSA to stabilize DNA interactions and minimize non-specific binding.

### Optical trapping assays

Optical trapping experiments were performed on an acoustically isolated custom-built optical trap around the body of a Nikon Ti-E inverted microscope (Belyy et al., 2014). A 20 W, 1064 nm ytterbium linear polarized CW laser (IPG Photonics) was used to trap beads. The trapping laser was focused on the imaging plane with a 100 x 1.49 N.A. apochromat oil immersion objective (Nikon). Axonemes purified from sea urchin sperm were allowed to attach to the coverslip surface and imaged via brightfield microscopy on a monochrome CCD camera (The Imaging Source). These axonemes were brought to the center of the field of view with a locking XY stage (M-687, Physik Instrumente). The head position was tracked by imaging the back focal plane of a 1.4 N.A. oil immersion condenser (Nikon) on a position-sensitive diode (PSD, First Sensor). The PSD response was calibrated by raster scanning trapped beads using a pair of acousto-optical deflectors (AODs, AA Optoelectronic). The power of the trapping laser was adjusted with a half-wave plate mounted on a motorized rotary mount (New Focus). The trap stiffness was then calculated by fitting the power spectrum of the trapped bead to a Lorentzian. Trapped beads were lowered onto the axoneme surface using a piezo flexure objective scanner (P-721, PIFOC, Physik Instrumente). Stall force measurements were performed against a stationary trap where motors were allowed to pull beads away from the trap center until detachment. In fixed trap assays, typical trap stiffnesses were 0.08-0.10 pN/nm for single motor, 0.1 pN/nm for two motors, and 0.2 pN/nm for three motors. Force feedback measurements were performed at a constant force by adjusting the position of the trap center with the AOD to maintain a fixed separation of the bead from the trap center, 100 nm in all cases except 1 pN hindering and 0.5 pN assisting, which used 50 nm. The desired force was achieved by tuning the trap stiffness accordingly, and the trap position was updated at 100 Hz. In all measurements, the bead position was acquired at 5 kHz, and the motor concentration was reduced until less than 30% of beads demonstrated motility within 100 s to ensure more than 90% of events were driven by a single motor.

### Motility assays

Motility assays were performed using surface-immobilized axonemes or microtubules. Axonemes were diluted 1:10 in MB and added directly to an untreated flow chamber for 3 min. For microtubule immobilization, flow chambers were assembled using coverslips functionalized with biotin-PEG-SVA (Laysan Bio). 20 μl of 1 mg mL^-1^ streptavidin was then added to the flow chamber for 1 min and washed with 30 μl MB. Biotinylated microtubules were then flowed into the flow chamber for 3 min. In both cases, the chamber was washed with 50 μl MBC (additionally supplemented with 0.5% pluronic and 10 μM taxol for microtubules). K560-GFP-SNAP was diluted to 1 nM in SB and flowed into a chamber containing axonemes. To test the oligo labeling of KIF5B, labeled KIF5B-GFP-SNAP was first incubated with a 10-fold excess of complementary Cy5-labeled oligo for 10 min before being diluted to 1 nM in SB and flowed into a chamber together with 10 nM MAP7. The ability of the long DNA handle to bind to streptavidin beads was tested with quantum dots. 2 μl of 100 nM DNA handle was incubated with 1 μl of 600 nM KIF5B and 0.5 μl of 1 μM Qdot 655 streptavidin-coated quantum dots (Invitrogen) for 30 min before being diluted to 1 nM in SB and flowed into a chamber together with 50 nM MAP7.

Fluorescence imaging experiments, except those using quantum dots, were performed using a custom-built multicolor TIRF setup described previously (DeWitt et al., 2012) with an effective pixel size of 160 nm after magnification. GFP and Cy5 were excited with fiber-coupled 488-nm (Coherent) and 633-nm (Melles Griot) laser beams through a Nikon TIRF Illuminator, and their fluorescent emissions were filtered through a notch dichroic filter and 525/40 and 655/40 bandpass emission filters (Semrock), respectively. Multicolor movies were collected using the time-sharing mode in MicroManager. Motility assays involving quantum dots were performed with a TIRF-aligned 488 nm laser beam (Coherent). Fluorescent emissions from the quantum dots were filtered through a 655/40 bandpass emission filter. Movies were collected on an ORCA Flash 2.0 CMOS camera (Hamamatsu) with an effective pixel size of 43.3 nm.

## Data Analysis

Motility assays were analyzed as described previously (Zhao et al., 2023). Optical trapping data were first converted from raw PSD voltage to either force or displacement. Stalls were defined as near-zero velocity of a bead with a minimum duration of 150 ms under hindering forces above 2.5 pN, followed by a snap-back of the bead to the trap center. Custom MATLAB software was used to identify stalls, and the stall force was defined as the average force exhibited in the terminal 20% of a stalling event, while the stall time was calculated as the duration in which the force exceeded 80% of the stall force. Peak force was defined as the four-point average of the highest force in each processive run. Only peak forces above 1.5 pN (corresponding to a 15-19 nm bead displacement) were analyzed to clearly distinguish runs from the tracking noise. Stall lifetimes were calculated by fitting the 1-CDF of stall durations to a double exponential decay. For multi-motor experiments, traces frequently exhibited rapid sequential detachments (Fig. 3C,E). The stall force for these measurements was taken to be the highest force sustained for at least 20 ms.

To identify potential runs under force feedback conditions, custom MATLAB software was used to identify all regions where the trap position is non-zero. The software then finds the subset of this region where force feedback is engaged. To exclude slips in velocity and run length calculations, a simple sliding window method is used to detect slips by measuring spikes in the standard deviation and then subsequently eliminating them. Runs with times lower than 100 ms, as well as runs with a velocity greater than 100 nm s^-1^ away from the next closest velocity, were excluded from analysis. The run length at each applied force was calculated by fitting the CDF of the end-to-end distances of the runs to one minus a single exponential decay function. Under 1 and 2 pN hindering forces, kinesin was occasionally able to reach the furthest displacement set for the trapping laser (1000 nm in the high z-force regime and 3000 nm in the low z-force regime). These instances were removed, and the maximum value of the CDF was reduced by a fraction exceeding this limit (Figure 2_figure supplement 2A). Velocities were calculated by dividing the end-to-end displacement of each run by the elapsed time and reported as the mean ± s.e. (Figure 2_figure supplement 2B). Detachment rate was calculated as the quotient of velocity divided by run length for all forces with positive run lengths. For force regimes without net forward motility, the detachment rate was calculated as the inverse of the time constant extracted from single exponential decay fits of the 1-CDF of the attachment time (Figure 2_figure supplement 2C).

For multi-motor experiments on a DNA chassis, occasional stalls exceeding the product of the expected motor copy number multiplied by 10 pN were excluded from analysis, as these were attributed to the beads carried by multiple chassis. An instantaneous >2 pN decrease in force to a non-zero force was categorized as a slip. The 1-CDF of the slips originating from forces above 8.1 pN and 10.8 pN (corresponding to 1.5x and 2x the stall force of a single kinesin) for the 2- and 3-motor chassis, respectively, were fit with a double exponential, and the time constant reported is a weighted average of the two time constants of the fit (Elshenawy et al., 2019).

## Statistical Analysis

The p-values were calculated by the two-tailed t-test in Prism and Origin. CDFs were calculated in MATLAB.

## Data Availability

The main data supporting the findings of this study are available within the article and its Supplementary Figures. The plasmids will be deposited at AddGene. Raw microscopy data will be made available by the corresponding authors upon request.

## Code Availability

The custom code used to analyze experimental data is uploaded to the Yildiz Lab code repository (www.yildizlab.org/code_repository) and GitHub (https://github.com/Yildiz-Lab/YFIESTA, doi: 10.5281/zenodo.8415156).

## Acknowledgements

The authors are grateful to V. Belyy and Y. Ezber for technical assistance with optical trapping and to the members of the Yildiz laboratory for helpful discussions. This work was supported by NIH (R35GM136414) and NSF (MCB-1055017) to A.Y.

## Author contributions

A.Y. conceived the project. A.M.H. developed the labeling scheme. A.Y. and A.M.H. designed the experiments. A.M.H. performed all experiments. A.Y. and A.M.H. wrote the paper.

## Competing interests

The authors declare no competing interests.

---

**Figure 1_figure supplement 1.**
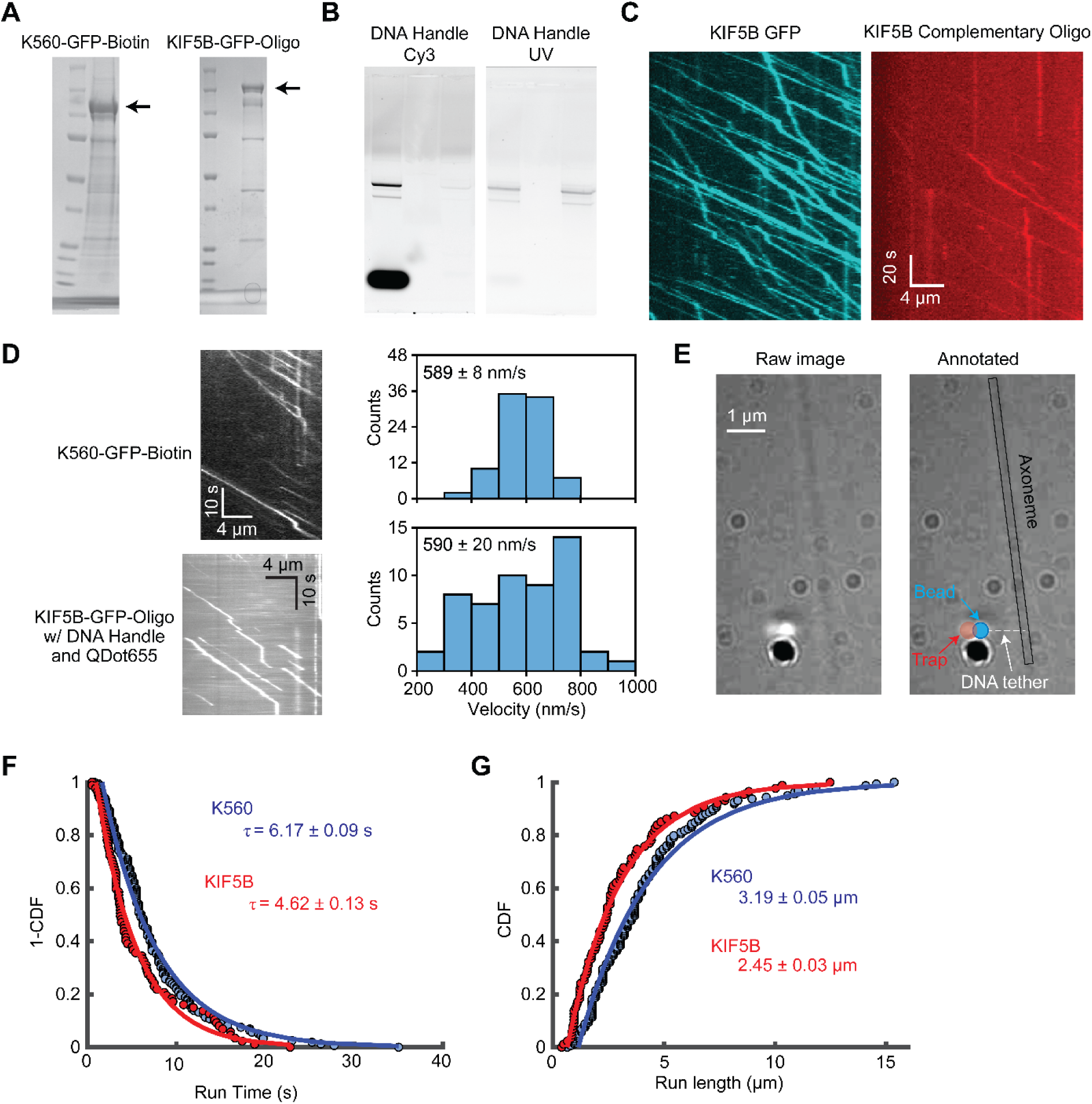
Purification of kinesin and DNA handle with experimental controls. **A)** SDS-Page denaturing gel of constructs used. K560-GFP-SNAP was labeled with biotin, and KIF5B-GFP-SNAP was labeled with a BG-functionalized DNA oligo before eluting the motor from the IgG beads during purification. **B)** The long DNA handle was run on a 0.8% TAE agarose gel. Left: Handle incubated with a 20-fold excess of complementary Cy3 oligo and imaged in the Cy3 channel of a Typhoon FLA 9500 fluorescence imager. Right: The same gel imaged under UV after staining in 2x GelRed for 40 minutes. **C)** Full-length KIF5B-GFP-SNAP was incubated with a 10-fold excess of complementary Cy5-labeled oligo and run in a motility assay. The assay was performed in the presence of 10 nM MAP7. **D)** Kymographs of K560-GFP-biotin (top, N = 89) and Qdot 655 streptavidin quantum dots being transported by oligo-labeled KIF5B attached to the long DNA handle (bottom, N = 53). Histograms of the respective velocities are shown to the right. **E)** Mechanical demonstration of a bead tethered to an axoneme by the long DNA handle. After tethering to an axoneme, the bead is pulled away from the axoneme by an optical trap. The bead moved freely for ∼1 µm and stayed in that position, demonstrating the formation of a long tether between the bead and the axoneme through the DNA handle. **F)** 1-CDF kinesin run time under unloaded conditions in single-molecule fluorescence imaging assays (N = 160 for K560 and 88 for KIF5B). Solid curves represent a fit to a single exponential decay function to calculate the lifetime of processive runs (τ, ±S.E.). **G)** CDF of kinesin run length under unloaded conditions in single-molecule fluorescence imaging assays. Solid curves represent a fit to a single exponential decay function to calculate the mean run length (±S.E.). In D, F, and G, KIF5B assays were performed in the presence of 50 nM MAP7.

**Figure 2_figure supplement 1.**
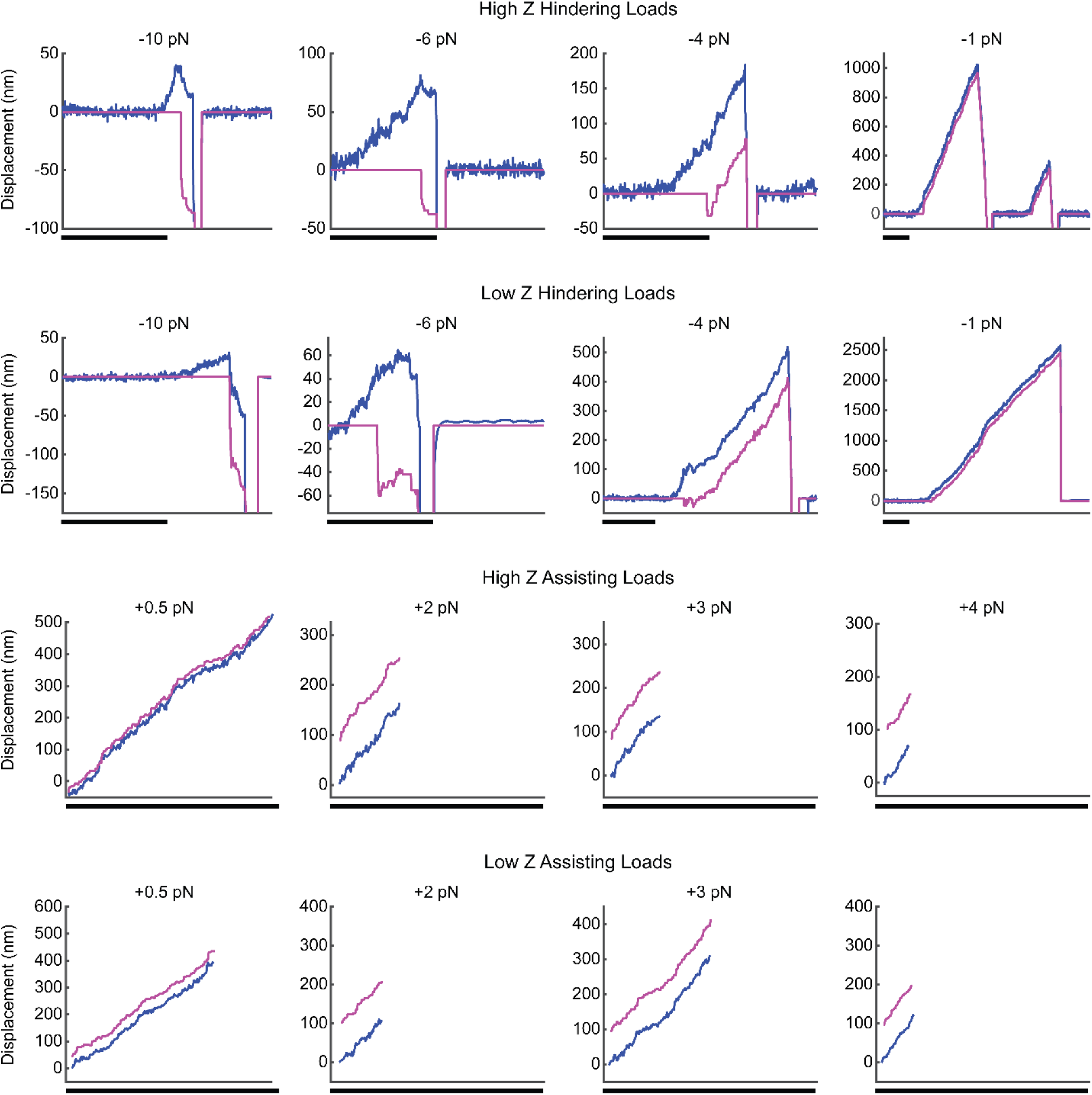
Example traces for force-feedback controlled trapping of kinesin with or without a DNA handle. Example traces for force feedback measurements under the high z-force and low z-force conditions in hindering and assisting directions. Scale bars are 1 s.

**Figure 2_figure supplement 2.**
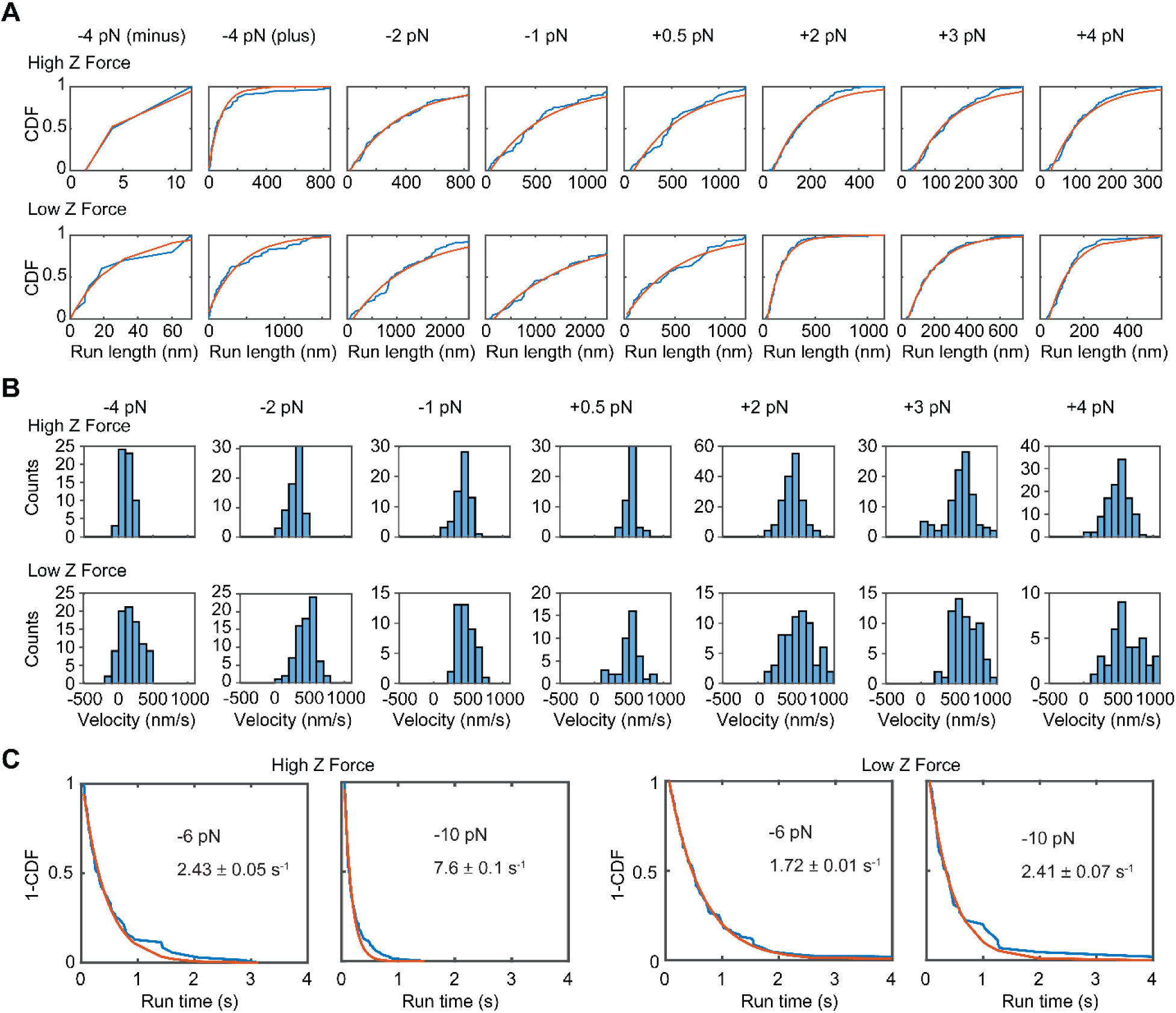
Raw histograms and fits to force-feedback controlled trapping of kinesin with or without a DNA handle. **A)** CDFs of run length at each condition (blue) and the corresponding fit (red). Under the 4 pN hindering condition, a small number of events demonstrated net negative run lengths. A weighted average was used to calculate the run length shown in Fig. 2. From left to right, N = 3, 55, 69, 65, 50, 167, 100, 115 on the top row, and 11, 78, 76, 44, 42, 65, 32, 39 in the bottom row. **B)** Histograms of velocities for each condition. From left to right, N = 60, 69, 65, 50, 167, 100, 115 in the top row and 89, 76, 44, 42, 65, 32, 39 in the bottom row. **C)** 1-CDFs of motor attachment time to the microtubule without net positive displacement under stall or superstall forces. Solid red curves represent a fit to a single exponential decay to calculate the detachment rate (From left to right, N = 167, 133, 73, 43).

**Figure 3_figure supplement 1.**
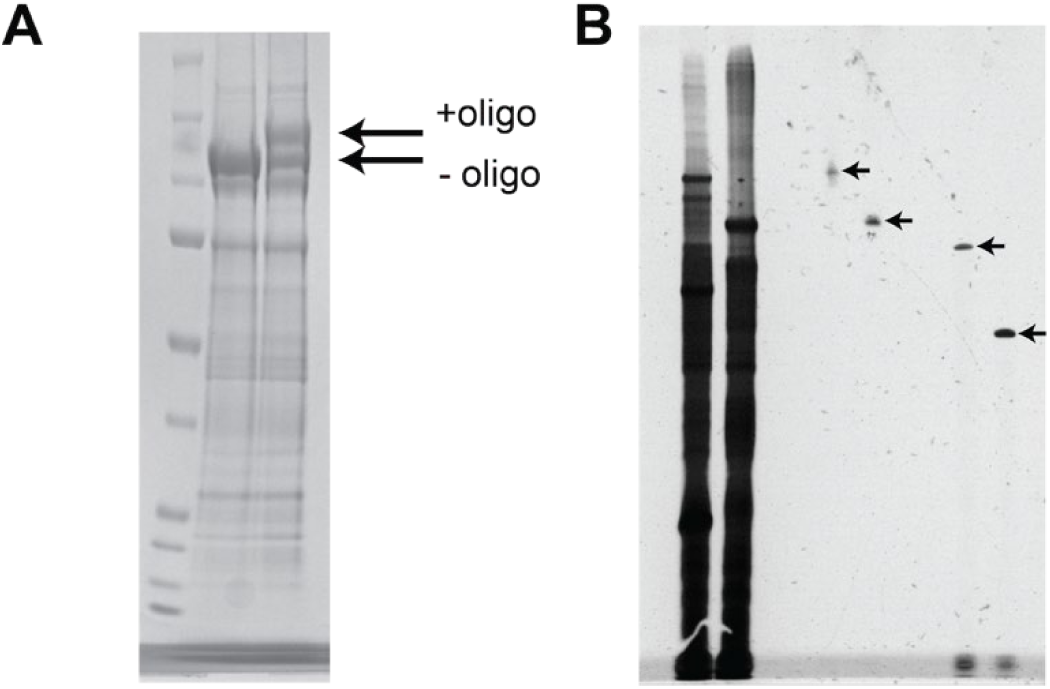
Purification and stall force of multi-motor chassis. **A)** SDS-PAGE denaturing gel to quantify oligo labeling of K560-GFP-SNAP. Left lane: molecular weight marker. Middle lane: K560-GFP-SNAP. Right Lane: K560-GFP-SNAP labeled with DNA oligo. Arrows show that the oligo-labeled motor is discernible from the unlabeled motor on the gel. **B)** Gel extraction of multi-motor chassis. The left two lanes show 3- and 2-motor chassis for high z-force measurements before gel extraction. The middle two lanes show 3- and 2-motor chassis for high z-force measurements after gel extraction. The right two lanes show 3- and 2-motor chassis for low z-force measurements after gel extraction.

**Figure 3_figure supplement 2.**
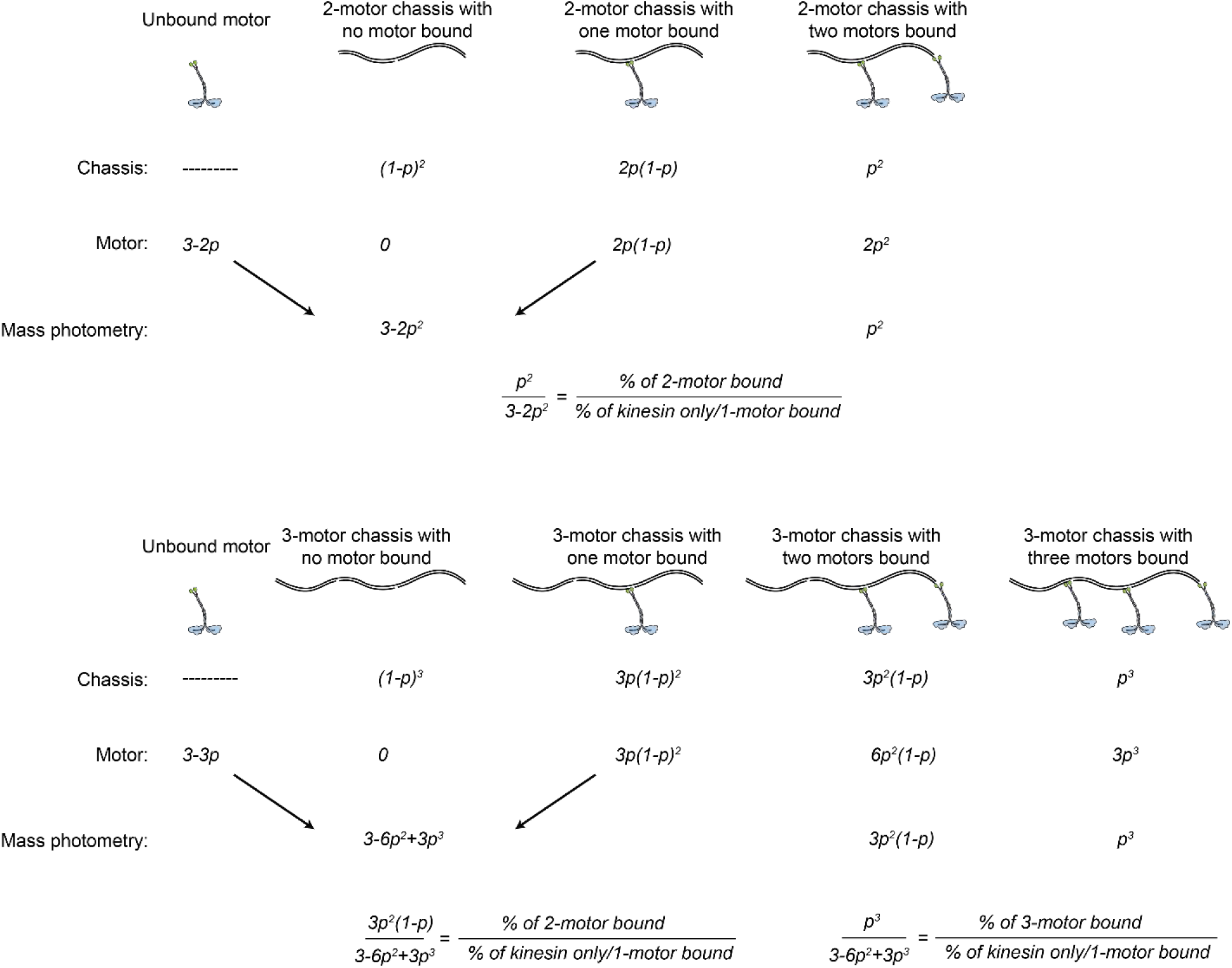
Model for estimating the percentage of DNA chassis bound to two- or three-motors in mass photometry. Unbound motor is 3-fold excess of the DNA chassis. *p* is the probability of binding of kinesin to the DNA chassis. The model assumes no cooperativity between the binding sites on the chassis. Motor bound to chassis was calculated from the probability of chassis multiplied by the number of motors bound to the chassis. Mass photometry cannot distinguish an unbound motor and chassis with one motor bound. *p-*values were calculated from the ratios of the percentages of distinct mass populations detected by mass photometry.

**Figure 3_figure supplement 3.**
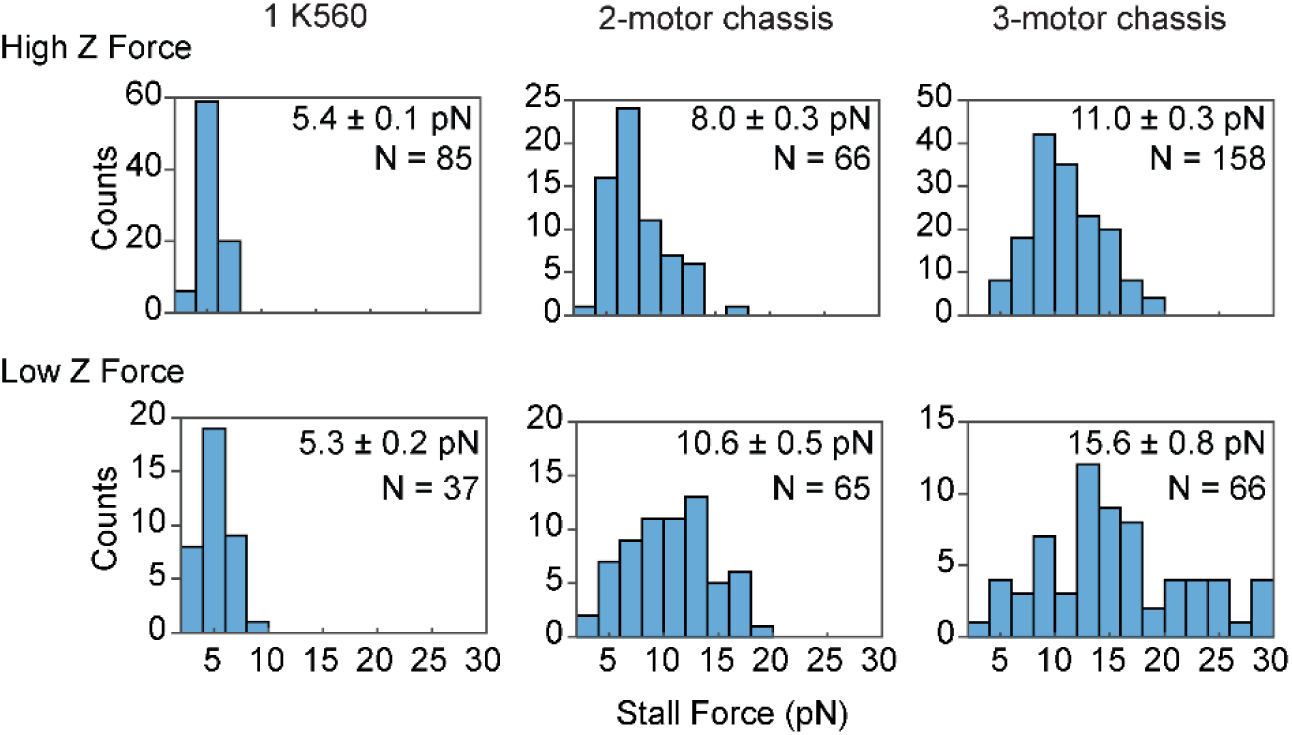
Stall force histograms (mean ± s.e.) of K560, and 2- or 3-motor chassis under high and low z-force conditions.

## Notes

### Competing Interest Statement

The authors have declared no competing interest.

### Summary of Updates

In the revised manuscript, we carefully reported the statistics of experimental measurements, provided an explanation for the discrepancy between our results and DNA tensiometer assays, and added a supplemental figure (Figure 3_figure suplement 2) to show our calculation for the percentage of DNA chassis with multiple kinesins. We also made additional changes throughout the manuscript to improve the clarity of our work.

